# Residual photoreceptors affect the response of a degenerate retina to electrical stimulation

**DOI:** 10.64898/2025.12.15.694454

**Authors:** Keith Ly, Mohajeet B. Bhuckory, Davis Pham-Howard, Anna Kochnev Goldstein, Nathan Jensen, Daniel Palanker

## Abstract

Patients with age-related macular degeneration (AMD) implanted with the PRIMA photovoltaic subretinal prosthesis demonstrated letter acuity closely matching the device’s 100 µm pixel size. Improving visual acuity requires smaller pixels, which, in turn, require relaxation of electric field confinement to maintain effective stimulation of bipolar cells. Eliminating local return electrodes broadens the electric field but may inadvertently engage residual photoreceptors adjacent to the implant, thereby altering electrically evoked visual percepts. Here we quantify the contribution of residual photoreceptors to electrically evoked retinal responses across various implant configurations.

Monopolar photovoltaic arrays with 20 µm pixels and bipolar arrays with 100 µm pixels were implanted subretinally in Long Evans rats, resulting in local degeneration of photoreceptors directly above the device. Responses were compared with those obtained in RCS rats, which lack functional photoreceptors. Implants were activated by patterns of 880 nm laser at pulse durations varying from 0.5 to 10 ms. Visually evoked potentials were measured in scotopic and photopic conditions, with and without the intravitreal application of mGluR6 agonist L-AP4 to block photoreceptor-driven ON pathways. Experimental thresholds were interpreted using a computational model of retinal network activation in distinct electric field geometries.

In locally degenerate retina stimulated with monopolar arrays, blocking photoreceptor input yielded a rheobase (0.06 mW/mm²) and chronaxie (∼3 ms) of the stimulation threshold, matching that measured in fully degenerate RCS retina, indicating direct activation of bipolar cells. In contrast, when photoreceptor input was intact, stimulation thresholds decreased significantly, and dark adaptation further modulated the threshold in a pulse-duration dependent manner. Bipolar arrays provided identical thresholds in locally degenerate and fully degenerate retina (0.2 mW/mm²) when stimulation was confined to the implant center; however, shifting stimulation to the implant’s edge lowered the thresholds, revealing a contribution from adjacent photoreceptors.

These findings demonstrate that residual photoreceptors can substantially influence responses evoked by subretinal prostheses in a degenerate retina, with direct consequences for perceptual uniformity in patients, such as edge brightening. These results provide guidance for design of the next-generation higher-resolution implants, supporting the use of local return electrodes to maximize resolution while minimizing the effect of photoreceptors in clinical applications.

## 1. Introduction

Patients afflicted by retinal degenerative diseases, such as age-related macular degeneration (AMD) and retinitis pigmentosa (RP), experience loss of vision due to death of photoreceptors [1]. Retinal prostheses aim at restoring visual function in such patients by stimulating the remaining interneurons. In patients with AMD, photoreceptors in the central macula, which is responsible for high-resolution vision, are lost, but the inner retinal neurons are largely preserved. Subretinal prostheses provide central vision by stimulating these interneurons, whilst not affecting the peripheral healthy retina.

One such device, the PRIMA system, provides prosthetic vision to AMD patients with letter acuity up to 20/420, matching the sampling limit of its 100 µm pixels [2]. With the current bipolar design of photovoltaic arrays (Figure 1A, B), where active electrodes are surrounded by local returns in each pixel, reducing the pixel size decreases the field penetration into tissue, thus precluding retinal stimulation with much smaller pixels [3]. Monopolar arrays (Figure 1C, D) provide deeper penetration of electric field, enabling the use of smaller pixels, down to 20 µm, geometrically corresponding to 20/80 acuity in human eye [4].

**Figure 1.**
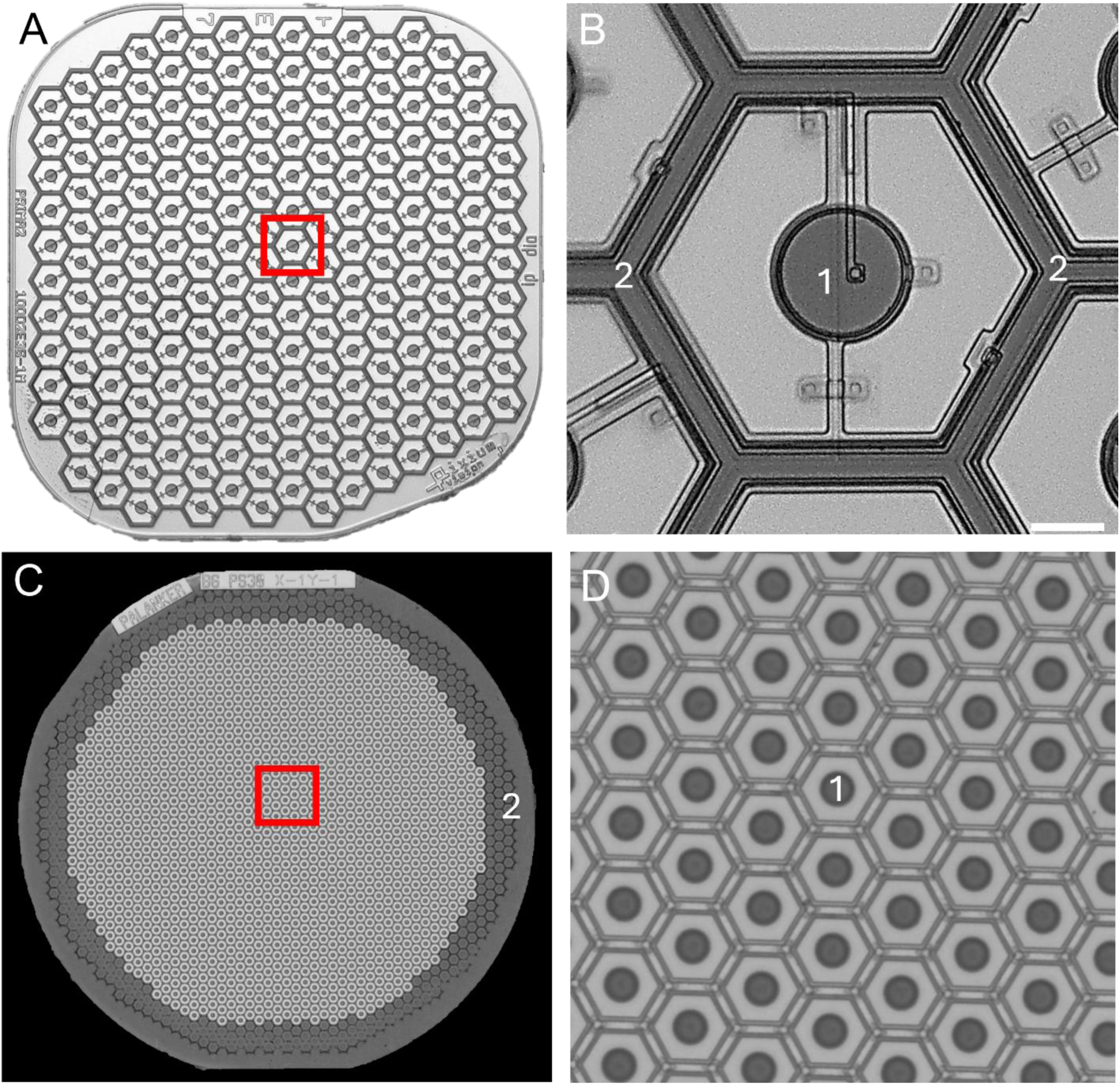
Photovoltaic implants used in this study. Implants are 1.5mm in width. **A.** Implant with bipolar electrode configuration. 100 µm pixels (**B**) with an active central electrode (1) and local returns (2) **C.** Monopolar implant configuration. 20 µm pixels (**D**) with active electrodes (1) and a global surrounding return electrode (2).

In this study, we explore the effect of the residual photoreceptors outside the implant on its electrical stimulation. For this purpose, we implanted Long-Evans (LE) rats subretinally with the monopolar and bipolar photovoltaic arrays, which resulted in degeneration of photoreceptors directly above the implant, while being preserved outside the device [5]. This condition resembles local retinal degeneration in AMD. To examine the role of photoreceptors in subretinal stimulation, we measured the stimulation thresholds with and without the mGluR6 agonist L-AP4, which blocks the ON pathway from photoreceptors – by far the prevailing pathway in the rod-dominant rat retina. For comparison, we performed similar measurements in RCS rats after a complete degeneration of all photoreceptors.

We also modelled both healthy and degenerate photoreceptors in the software NEURON to assess their effect on electrical stimulation above and around the devices of monopolar and bipolar configurations.

## 2. Methods

All experimental procedures were conducted in accordance with the Statement for the Use of Animals in Ophthalmic and Vision research of the Association for Research in Vision and Ophthalmology (ARVO) and approved by the Stanford Administrative Panel on Laboratory Animal Care (APLAC). Animals were maintained at the Stanford Animal Facility under 12 h light/12 h dark cycles with food and water *ad libitum*. Animals that required dark adaptation were placed in complete darkness for at least 12 hours.

### 2.1. Surgical procedures

Animals were anesthetized with a mixture of ketamine (75 mg/kg) and xylazine (5 mg/kg) injected intraperitoneally. All rats were implanted with three transcranial screw electrodes: one over each hemisphere of the visual cortex (4 mm lateral from midline, 6 mm caudal to bregma), and a reference electrode (2 mm left of midline and 2 mm anterior to bregma). To ensure stable results, measurements were performed at least 8 weeks post implantation [6]. Bipolar photovoltaic devices with 100µm pixels and monopolar photovoltaic devices with 20µm pixels of 1.5 mm in diameter and 30 μm in thickness were implanted subretinally, as previously described [4, 7]. Retinal implants were placed in the temporal-dorsal region, approximately 1 mm away from the optic nerve. Royal College of Surgeons (RCS) rats were used after age P180, when photoreceptors completely disappeared [8]. In Long Evans (LE) rats, loss of the outer nuclear layer (ONL) (Figure 2) by 8 weeks was checked with optical coherence tomography (OCT; HRA2-Spectralis; Heidelberg Engineering, Heidelberg, Germany). Histology slices also demonstrate that photoreceptors are not present above the implant.

**Figure 2.**
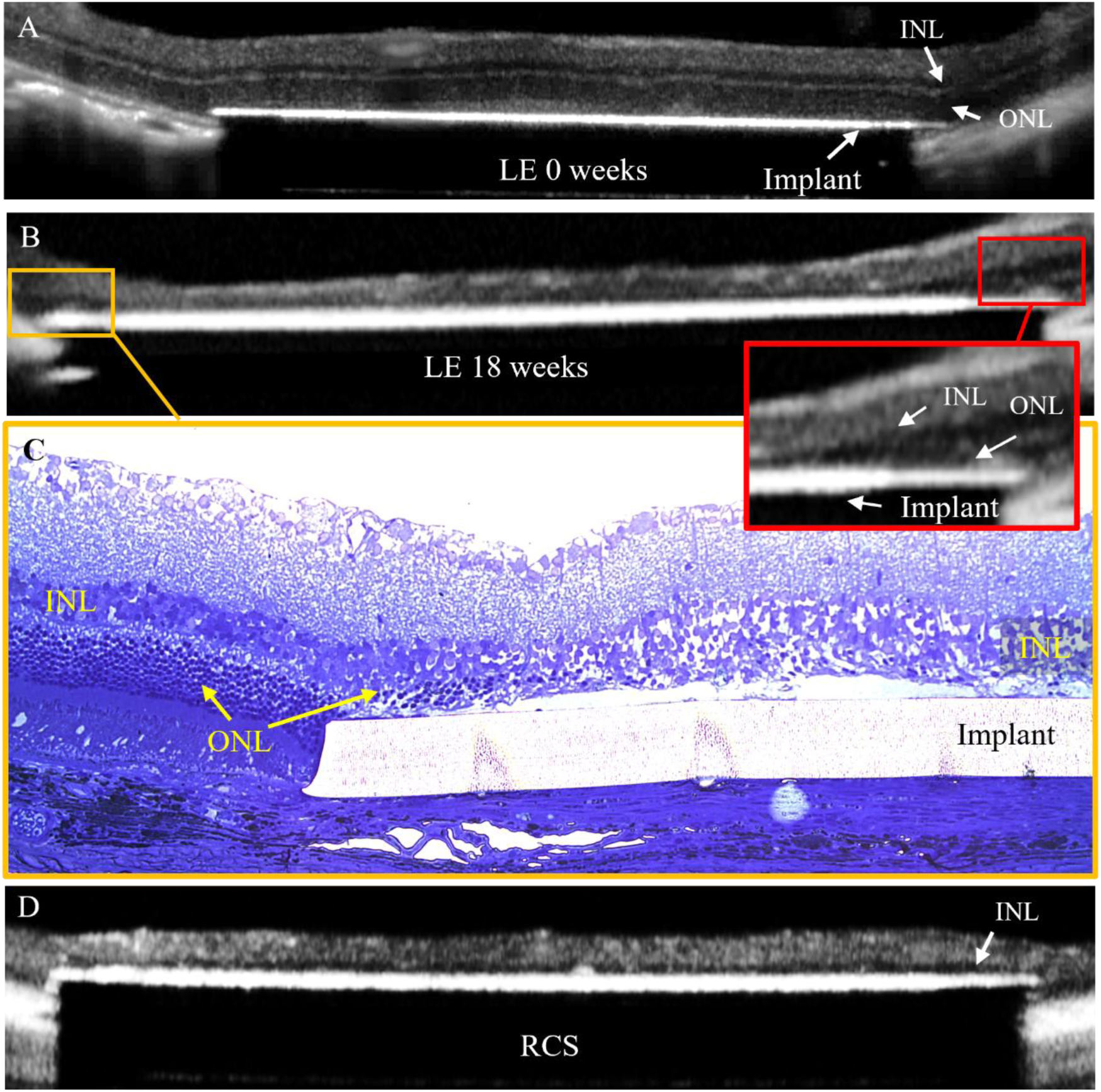
**A.** OCT B-scans of the subretinal implant in LE rats at day 0. **B.** After 18 weeks post-op, the ONL above the implant completely disappears after 8 weeks. **C.** However, there are still remnant photoreceptors present above the edge of the implant, as shown in the histology slice. **D.** OCT B-scan of the subretinal implant in an RCS rat.

To check efficiency of L-AP4 (0.8 mM) in blocking the photoreceptors, visually evoked potential (VEP) response to full-field green light flashes was recorded in LE rats before and after the intravitreal injection in a control area 1.5 mm nasal from the implant, where photoreceptors are present. The VEP signal was lost and did not reappear for 3 hours post injection of the blocker. Additionally, this was also checked for the full cocktail of blockers (0.8 mM L-AP7, 0.8 mM L-AP4, 0.2 mM NBQX, 0.4 mM strychnine, and 0.32 mM picrotoxin) and the signal also did not reappear for 3 hours post injection. For day-0 VEP measurements, implantations, and injections were conducted under red light to ensure that the retina was dark-adapted prior to recordings.

### 2.2. Electrophysiological measurements

VEPs were recorded using the Espion E3 system (Diagnosys LLC, Lowell, MA) at a sampling rate of 4 kHz and averaged over 250 trials. For degenerate conditions, VEP measurements were conducted at least 16 weeks after subretinal implantation to ensure stabilization of the VEP amplitudes [6] and complete degeneration of photoreceptors above the implant [5, 7]. VEP experiments with degenerate retinae were conducted in photopic conditions (room lighting) using experimental setup as previously described [3, 4]. For stimulation of photoreceptors above the implant, VEP measurements were performed right after implantation (day-0), when photoreceptors are still present. For experiments in scotopic conditions, animals were dark adapted for 12 hours.

Typical VEP waveforms are shown in Figure 3A. Peak-to-peak VEP amplitude was calculated as a difference between the first trough in the signal within 120 ms of the stimulus onset and the peak within 100 ms after the trough (Figure 3A). Noise was determined in a similar fashion, 320 ms after the stimulus onset. Stimulation threshold was defined as the irradiance required to obtain a signal-to-noise ratio (SNR) of 1 through linear extrapolation of 2 points with an SNR > 1 (Figure 3B). The threshold was checked by ensuring that at irradiance below the extrapolated threshold, no signal was detected (SNR<1). Error from linear extrapolation of the lowest 2 points with an SNR > 1 was less than 15% in comparison to using 10 points. To determine the strength-duration curve of retinal stimulation, the thresholds *I(t)* were measured at pulse durations *t* ranging from 0.5 to 10 ms and fitted with the Weiss equation: 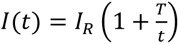, where *I_R_* is a rheobase and *T* – is a chronaxie [9] (Figure 3C, D). The VEP data were analyzed using custom code in MATLAB.

**Figure 3.**
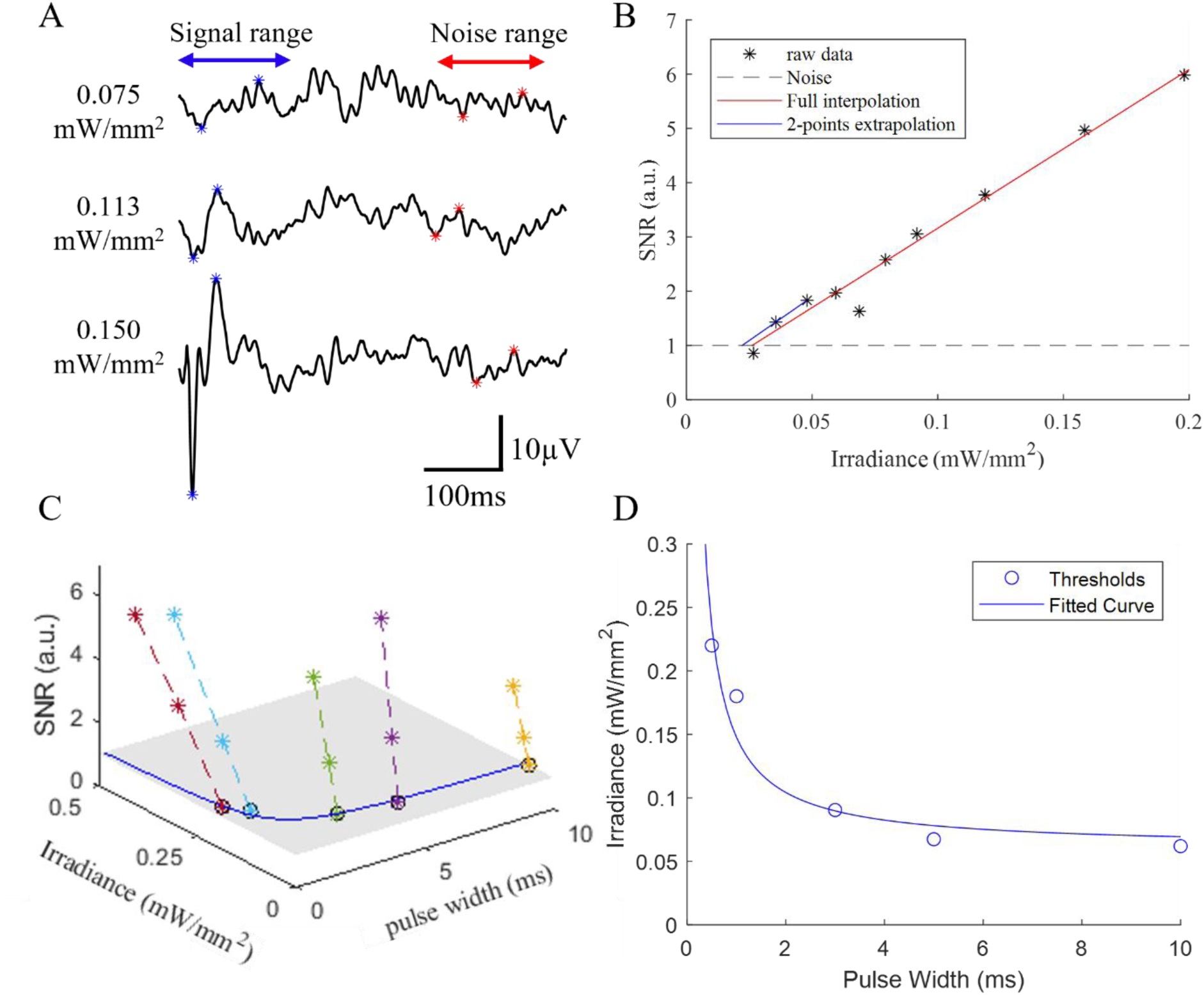
**A.** Example VEP waveforms demonstrating an increase in VEP signal with increasing irradiance. The signal is shown as blue asterisks (*) whilst the noise is shown as red asterisks (*). **B.** Extrapolation of the threshold (blue) is calculated by taking two points above the noise level. The signal-to-noise ratio (SNR) scales linearly with irradiance close to the threshold (red). **C.** Extrapolation of the strength duration (S-D) curve based on thresholds at multiple pulse durations. **D.** An example of an S-D curve fitted to Weiss’ equation.

### 2.3. Modelling of neural stimulation

Electric field in the retina was calculated using Retinal Prosthesis Simulator (RPSim) - a software package that models electric current generated by a photovoltaic array under patterned illumination and calculates electric potential in electrolyte by combining a finite-element method-based physics engine (COMSOL) with an electrical circuit solver [10, 11]. Here, a degenerate rat retina was modelled as a 126 µm thick layer of 1000 Ohm*cm resistivity [10], with the microelectrode array placed immediately below the inner nuclear layer (while for healthy retina, placed below the PR outer segments). Thickness of healthy rat retina was assumed 160 µm based on OCT imaging. Stimuli consisted of a single rectangular pulse with magnitudes defined by RPSim, reproducing the experimental conditions.

Healthy photoreceptors were modelled as 70 µm long cells which includes a soma diameter of 8 µm to account for the soma, inner segment, and outer segment. Partially degenerated photoreceptors were modelled as containing only the soma, which corresponds to the ONL observed at the edge of the implant in OCT (external limiting membrane is not visible), estimated to be 20 µm in length. Ion channels of the photoreceptor were based on previously published models [12, 13], modified to include multiple compartments (35 for healthy, 10 for degenerate) to improve sensitivity to extracellular electric fields. To probe the response to electric field, photoreceptors were placed between 500 µm and 1000 µm from the center of the implant at 20-25 µm increments.

## 3. Results

### 3.1. Stimulation thresholds in LE and RCS rats

Dependence of the stimulation threshold on pulse duration with monopolar array in LE rats under photopic conditions (n = 4) is shown in Figure 4A (black). With 10 ms pulses, stimulation threshold was 0.024 ± 0.007 mW/mm^2^, which is nearly 3 times lower than the 0.060 mW/mm^2^ measured in RCS rats (figure 4A, magenta) [4]. The 3.2 ms chronaxie of the Weiss equation-fitted Strength-Duration (S-D) curve indicates activation of the bipolar cells [14, 15], like in RCS rats [3]. The 0.024 mW/mm^2^ irradiance corresponds to a voltage drop of about 4 mV (obtained from modeled electric fields) across bipolar cells – much lower than in our previous studies in RCS rats, where the threshold at 10 ms was approximately 11 mV [3].

**Figure 4.**
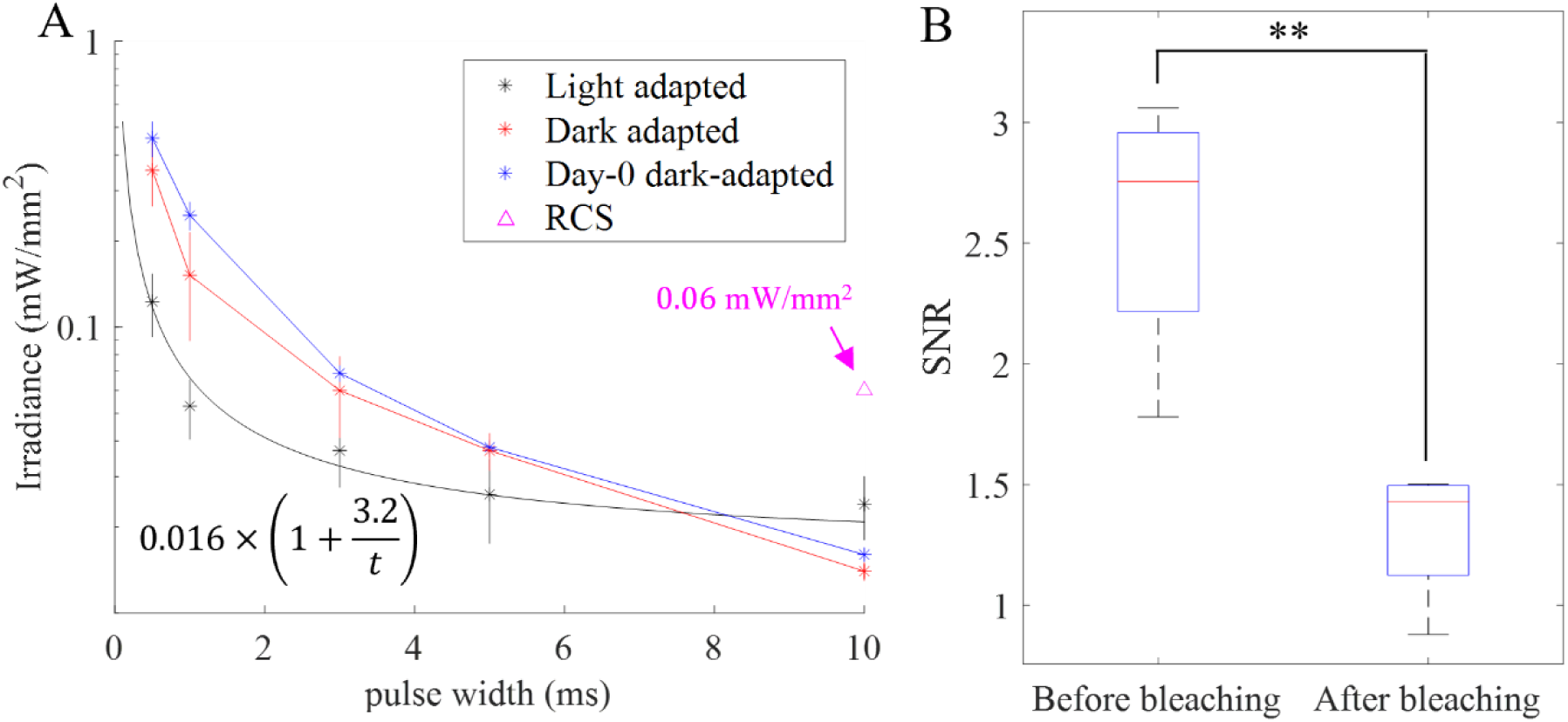
**A.** Strength duration (S-D) curves for monopolar implant with 20 µm pixels: irradiance thresholds as a function of pulse duration in LE rats >16 weeks post-op unless otherwise stated, under different lighting conditions. Under room lighting, the rheobase is 0.016 mW/mm^2^ (black), much lower than previously published [4, 6] with RCS rats (magenta). Dark adaptation changes the S-D curve, revealing direct stimulation of photoreceptors (red). S-D curves at day-0 match S-D curves from fully degenerated retina above the implant (blue). As PRs are fully depolarized in the dark, PRs directly above the implant do not respond to depolarizing electric field. Only PRs above the common return electrode where there is hyperpolarizing electric field can respond, indicating that the reduced threshold at longer pulse widths in dark-adapted degenerated retina occurs due to direct stimulation of PRs on the edge. **B.** Bleaching PRs with 20,000 lux of white light for 3 mins reduces the VEP signal (n = 4, p-value = 0.0069) of implant stimulation (0.2 mW/mm^2^), from a mean of 2.59 ± 0.49 before bleaching to 1.31 ± 0.25 after bleaching.

These results suggest that the threshold of bipolar cells may be reduced due to input from adjacent photoreceptors. Additionally, negative potential of the common return electrode along the edge of a monopolar implant may hyperpolarize the adjacent photoreceptors (Figure 2C) and thereby elicit a VEP.

To check the latter, we dark-adapted the animals to increase the photoreceptors sensitivity. In the dark, the neurotransmitter release rate of fully depolarized photoreceptors (at around -40mV) reaches its maximum, so any hyperpolarization results in a graded reduction of the glutamate release [16, 17]. The rod-dominated rat retina in dark-adapted conditions can respond to photon flux as low as 1 photon per second, which corresponds to hyperpolarization by about 1 mV [18, 19]. Dark adaptation significantly changed the strength-duration curve (Figure 4A, red). At 10 ms, the threshold decreased to 0.014 ± 0.001 mW/mm^2^, while at shorter pulses it increased. This new S-D curve did not fit the Weiss’ equation typical for other neurons [15], suggesting that different types of cells are activated at different pulse durations. We observed similar behavior in earlier ex-vivo studies using patch clamp recording [15], where photoreceptors dominated the retinal response at pulse durations exceeding 10 ms. This trend is also consistent with previous studies demonstrating that high frequency sinusoidal electrical stimuli (25 and 100 Hz) activate bipolar and ganglion cells, while lower frequency stimuli (5 and 10 Hz) activate photoreceptors [20].

VEP measurements under dark-adapted conditions immediately following implantation (day-0), when photoreceptors are still present above the implant, allow additional assessment of photoreceptor role in electrical response. Fully depolarized in the dark, photoreceptor terminals above the implant cannot respond to positive electric field, whilst in the negative field, photoreceptor terminals would be hyperpolarized, resulting in responses. The resulting S-D curve (Figure 4A, blue) was remarkably similar (R^2^ = 0.89) to the one recorded 16 weeks post implantation (Figure 4A, red), when photoreceptors above the implant are gone. This result indicates that dark adapted photoreceptors above the implant exposed to anodic subretinal pulses do not contribute, and that the response in the dark is driven by photoreceptors at the edges of the implant, experiencing a negative electric potential.

Interestingly, bleaching the photoreceptors with 20,000 lux of white light for 3 minutes reduced the amplitude of VEP above the threshold (at 0.2mW/mm^2^ at 10 ms, Figure 4B) - from the mean SNR of 2.59 ± 0.49 to 1.31 ± 0.25. This observation is consistent with our previous studies of the interactions between the peripheral retina and central prosthetic vision [21, 22]. Remarkably, these observations demonstrate two opposite effects of adjacent photoreceptors on electrical stimulation of bipolar cells: at room lighting, they reduced the electrical stimulation threshold compared to the degenerate retina, and yet after strong bleaching, they attenuate the magnitude of electrical response.

With bipolar pixel arrays (PRIMA) [23] illuminated in the center (Figure 5), stimulation threshold was much higher than with monopolar pixel arrays – 0.2 mW/mm^2^ at 10 ms pulse duration for both the LE (Figure 5, black) and in RCS rats (Figure 5, magenta) [3]. Under dark adaptation, stimulation thresholds with bipolar pixels did not change compared to photopic conditions (Figure 5, red). This finding indicates that, unlike monopolar arrays that have strong negative potential along the return electrode at the edge of the implants (Figure 1C), bipolar implants with an evenly distributed mesh of return electrodes (Figure 1B) do not stimulate even dark-adapted photoreceptors. This also indicates that since the thresholds are the same in light and dark adapted conditions, PRs do not affect the stimulation thresholds of BCs like in monopolar pixel arrays.

**Figure 5.**
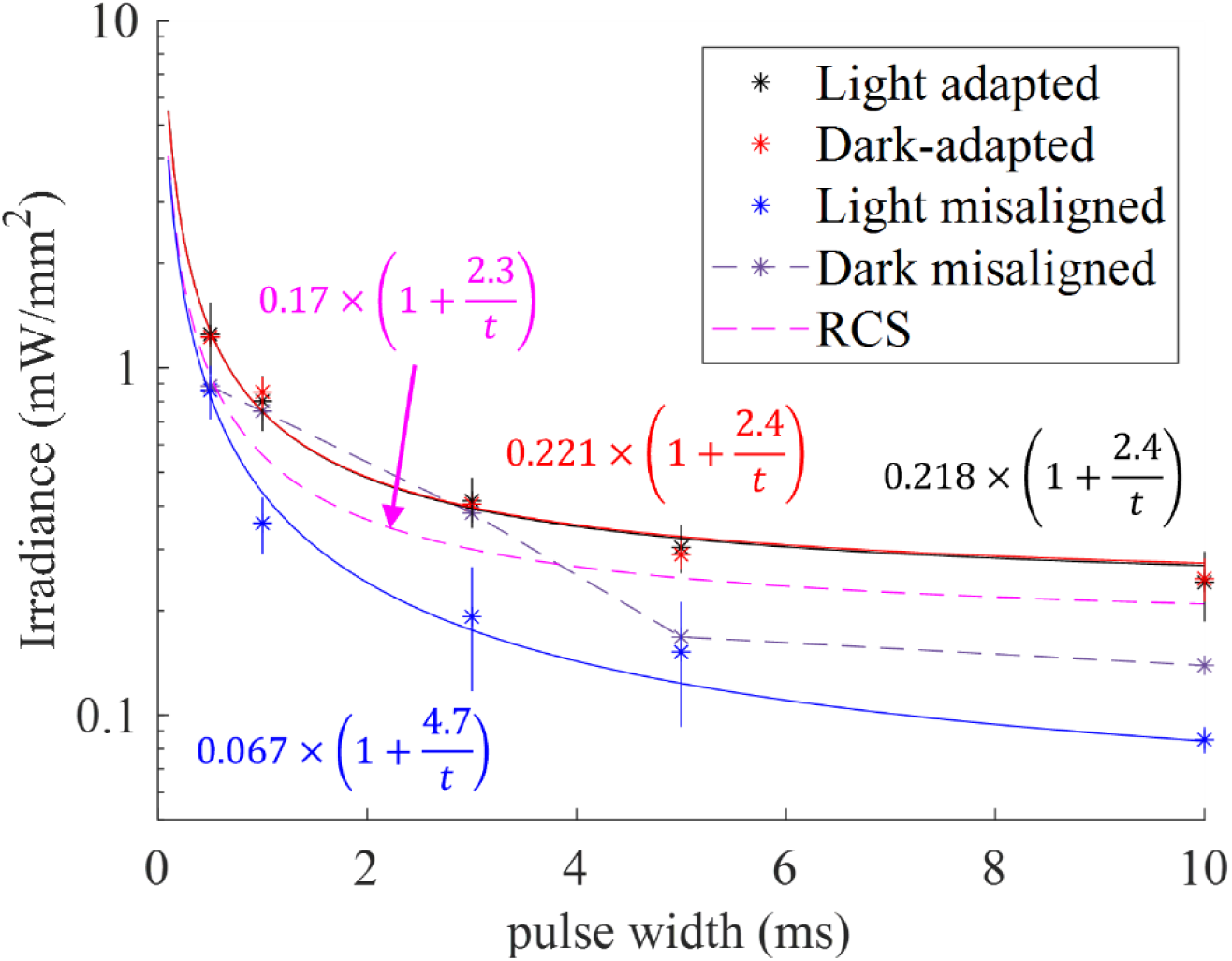
Strength duration (S-D) curves for PRIMA: irradiance thresholds as a function of pulse duration in LE rats >16 weeks post-op, under different lighting conditions. The S-D curves did not change between light (black) and dark-adapted (red) states, with a rheobase of 0.22 mW/mm^2^ and a chronaxie of 2.4 ms. Misaligning the beam to the edge of the PRIMA implant reduces the rheobase to 0.067 mW/mm^2^ (blue), by a factor of 3 from stimulation at the center (black). Dark-adapting the retina whilst misaligning the beam (purple, dashed) causes direct stimulation of photoreceptors at longer pulse widths (5 and 10 ms) but at a higher irradiance than in the light due to the lower negative potential from the local returns. Shorter pulse widths (0.5, 1, and 3 ms) cause BC stimulation. S-D curve for RCS rats with missing photoreceptors shown from previous publication [3] for comparison (magenta).

However, when the beam was shifted to the edge of the implant, the threshold in LE rats at room lighting decreased by about a factor of 3 – to 0.067 mW/mm^2^ (Figure 5, blue), revealing the influence of photoreceptors near the edge of the implant. In the dark, the threshold increased slightly (Figure 5, dashed line). Two effects can happen with the beam center shifted to the edge: (a) bipolar cells at the edge experience stronger electrical stimulation and their response becomes dominant in the population of cells above the implant. Photoreceptors under ambient lighting depolarize the ON BCs near the edge and reduce their stimulation threshold. (b) With a beam shifted to one side, stronger negative potential builds up on the dark part of the implant, and it starts stimulating the adjacent photoreceptors with longer pulses (Figure 5, dashed line), albeit not as strongly as with monopolar implants (Figure 4A, red line).

### 3.2. Effect of neurotransmitter blockers on stimulation

To further assess involvement of photoreceptors, we recorded VEP after intravitreal injection of L-AP4 (n = 3), the mGluR6 glutamate receptor agonist, which blocks the rod and ON cone inputs to bipolar cells by fully activating the glutamate receptors, which results in hyperpolarized ON bipolar cells (equivalent of dark conditions) [24, 25]. With monopolar implants, stimulation threshold at 10 ms increased to 0.067 ± 0.005 mW/mm^2^ (Figure 6A, green), closely matching the threshold in RCS rats: 0.060 mW/mm^2^ (Figure 6A, magenta) [4]. Fitting the data with Weiss equation yielded the chronaxie of 2.9 ms - in the range for bipolar cells stimulation (1.8 to 3.5 ms [3, 14]). This indicates that stimulation with the monopolar implant originates in bipolar cells, and after application of L-AP4, their resting potential is lower (and therefore the threshold is about 4 times higher) than when illuminated photoreceptors outside the implant provide a depolarizing input (Figure 6A – green vs. black line). Additionally, injection of the cocktail that blocks all synaptic transmission in the retina completely abolished the VEP signal at this irradiance, further indicating that the thresholds are driven by bipolar cells.

**Figure 6.**
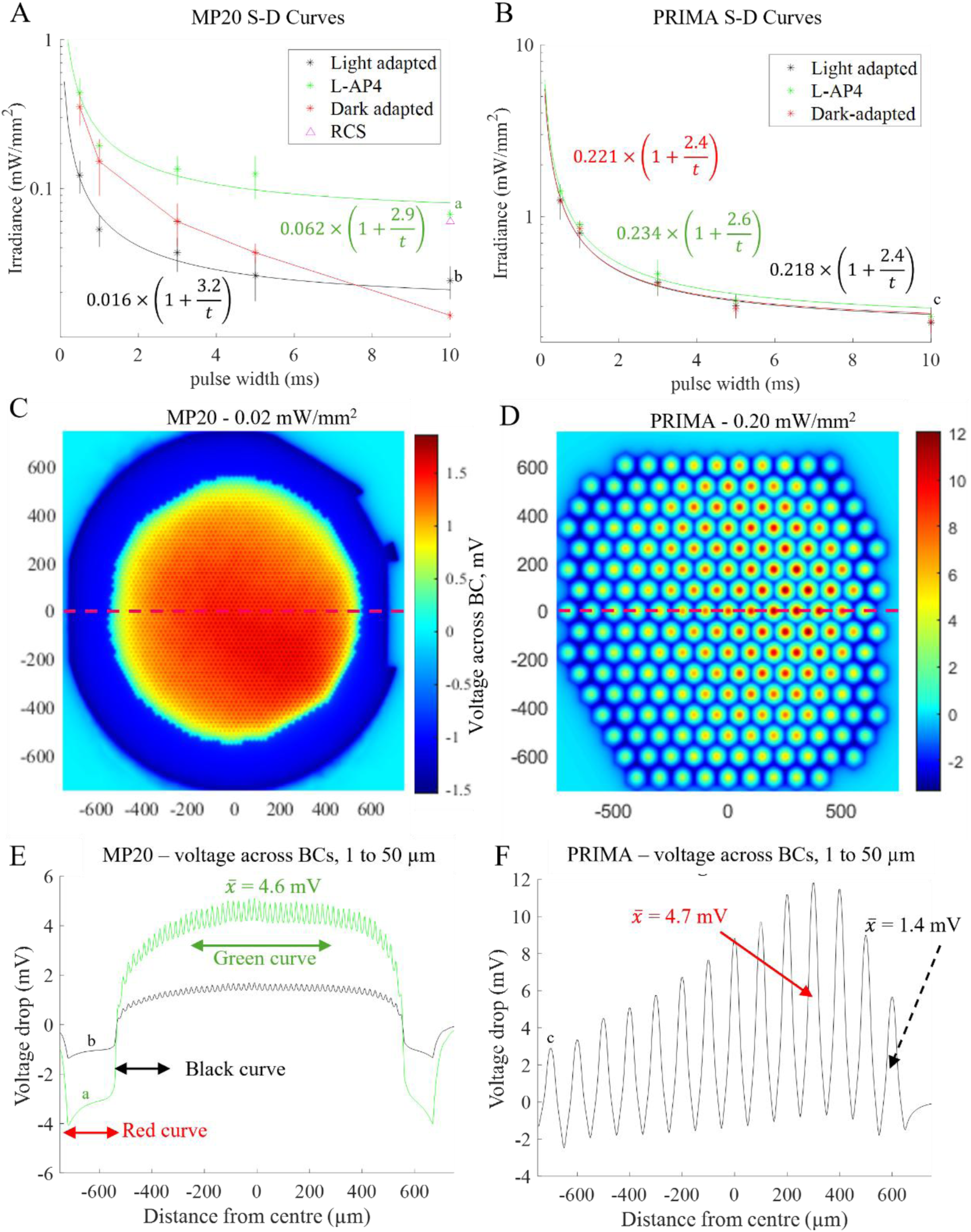
**A.** The rheobase of 20 µm monopolar pixel arrays at room lighting is 0.016 mW/mm^2^ (black). Dark-adaptation revealed stimulation of PRs (red). Intravitreal injection of L-AP4 (green) increased the threshold to 0.06 mW/mm^2^ matching the threshold previously recorded in RCS rats (magenta). **B.** With bipolar implants (PRIMA), thresholds with and without PR blocker L-AP4 are the same, suggesting that they are not affected by residual PRs. Dark-adaptation did not change the response (red). **C.** Electric field across bipolar cells at 0.02 mW/mm^2^ (50 µm) under monopolar configuration. Implant diameter is 1.5 mm. **D.** Electric field across bipolar cells at 0.20 mW/mm^2^ (50 µm) under bipolar configuration. The implant diameter is 2 mm. **E.** Voltage drop across BCs (from 0 µm and 50 µm) above the implant at 0.06 mW/mm^2^ (green, threshold at 10 ms in A, labelled “a”) and 0.02 mW/mm^2^ (black, threshold at 10 ms in A, labelled “b”). Average voltage drop across 100 µm is 4.6 mV. Arrows show regions responsible for the thresholds in A. **F.** Voltage drop across BCs (from 0 µm and 50 µm) above the implant at 0.20 mW/mm^2^ (black, threshold at 10 ms in B, labelled “c”) and the average across a 100 µm pixel (red*). Average voltage drop across 100 µm is 4.7 mV for a pixel in the center and 1.4 mV for a pixel on the edge.

Bipolar implants under room lighting exhibited similar thresholds in light and dark conditions (0.24 and 0.23 mW/mm^2^ at 10 ms, respectively), as well as with L-AP4 (0.22 mW/mm^2^, Figure 6B), indicating that, unlike with monopolar arrays, photoreceptors do not affect the bipolar cells threshold with bipolar devices in our illumination settings. This is likely due to non-uniform illumination in our setup which impacts the electric field from bipolar pixels, unlike monopolar arrays which create much more uniform field due to strong crosstalk between neighboring pixels, as shown in Figure 6 C-F. As shown in Figure 5 (blue vs. black line), the non-uniform illumination resulted in a stimulation threshold decrease by about a factor of 3 when we shifted the beam center to the edge of the bipolar pixels array. This measurement demonstrates that the stimulation threshold is defined by bipolar cells in the strongest field area, and when this area is close to the implants edge, illuminated photoreceptors outside the implant reduce the stimulation threshold by depolarizing the ON bipolar cells under room lighting conditions.

### 3.3. Modelling the photoreceptors and bipolar cells stimulation

To examine the electric potential drop across retinal neurons, we computed the electric field generated by photovoltaic arrays in electrolyte (1000 Ohm*cm resistivity) near the implant (Figure 6C-F). With monopolar arrays at stimulation threshold of 0.024 mW/mm^2^, the maximum voltage drop across bipolar cells (between 1 and 50 μm above the implant) was 1.7 mV, and with L-AP4 injection (0.067 mW/mm^2^) – 5.1 mV (Figure 6E). With bipolar arrays, there was no observable difference in the thresholds with and without L-AP4 (0.24 mW/mm^2^), corresponding to a maximum of 11.8 mV drop across bipolar cells. This is approximately 2.3 times that of monopolar arrays (Figure 6F). However, the average electric potential drop over the pixel area was 4.7 mV for both monopolar and bipolar arrays, indicating that VEP represents response of a population of cells, rather than just a few neurons localized in strongest field.

In LE rats, photoreceptors are preserved outside the implant, and cell bodies without the outer segments (OS) remain above implant near the edge (Figure 2B). To assess the photoreceptors’ response and the lateral range of retinal activation, we computed the electric field around the implant (Figure 7). Photoreceptors were placed at 20-25 µm increments between 500 µm and 1000 µm from the implant’s center (edge of the implant is at 750 µm).

**Figure 7.**
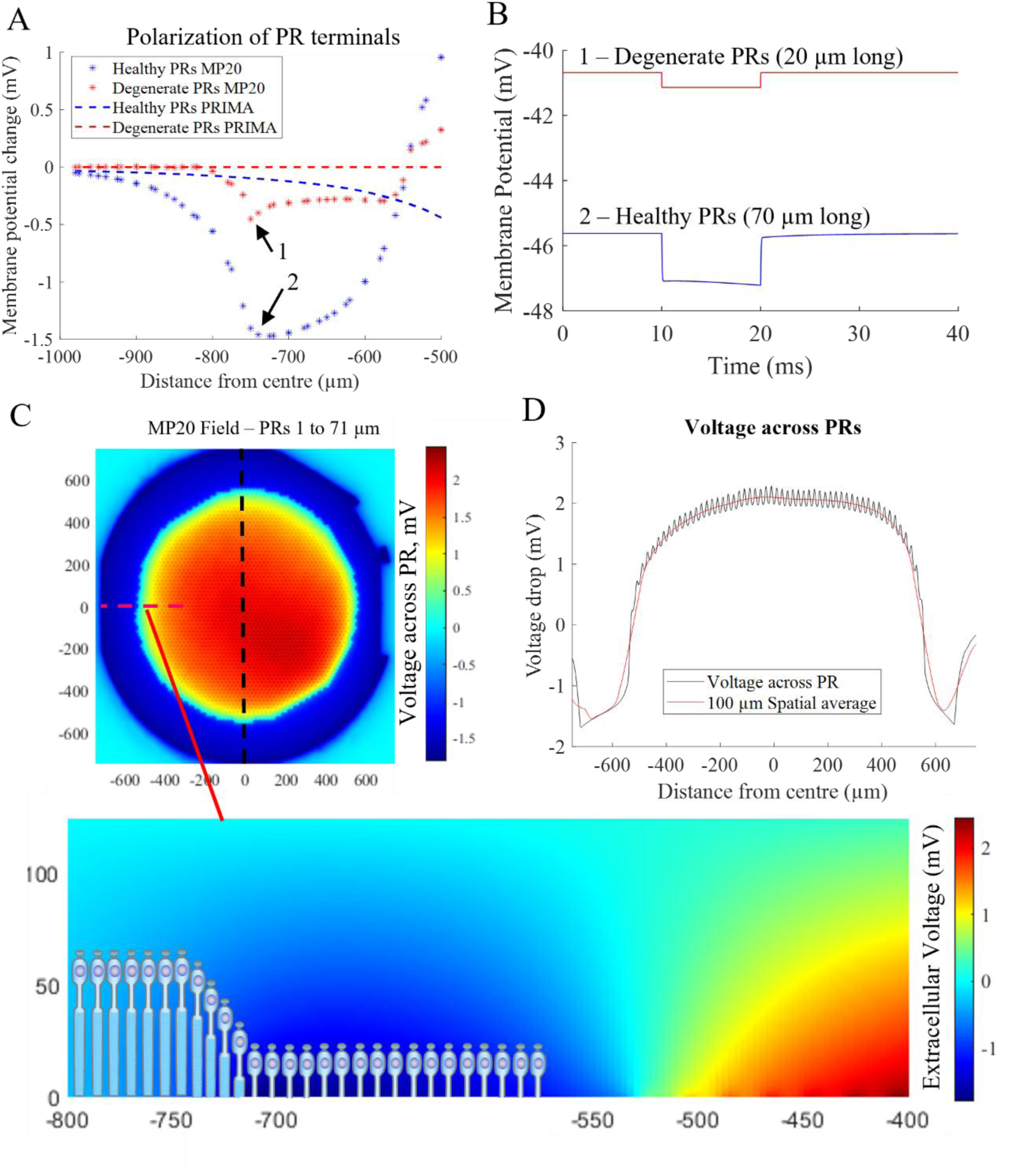
NEURON model of PR response under monopolar and bipolar electrode configuration. **A.** Maximum membrane potential change reveals thresholds for both degenerate PRs (0.5 mV hyperpolarization) and healthy PRs (1.5 mV hyperpolarization). Healthy PRs under bipolar electrode configuration are unable to reach threshold due to the constrained electric field from local returns. **B.** The response of both degenerate and healthy photoreceptors at the edge of the implant where the electric field change is the maximum. **C.** Electric field across healthy photoreceptors (70 µm) under monopolar configuration at 0.02 mW/mm^2^. Implant diameter is 1.5 mm. **D.** Voltage drop across PRs along the center of the implant (dashed line in C) demonstrating maximum hyperpolarization occurs on the edge of the implant. Average voltage across 5 pixels (100 µm) shown in red.

The maximum hyperpolarization of the photoreceptor terminals occurred within the negative potential zone at the edge of the monopolar implant (Figure 7A). At stimulation threshold of 0.020 mW/mm^2^, this was -1.5mV in healthy photoreceptors (day 0) above the implant (Figure 7B, blue), and by -0.45 mV in degenerate photoreceptors (Figure 7B, red). This is within the range of hyperpolarization of dark adapted rod photoreceptors in response to single photons per second: -0.64 ± 0.48 mV [19]. In the area of positive potential above the central part of the implant, photoreceptor terminals were depolarized (Figure 7C). Since at day-0 the animals were dark-adapted, photoreceptors were fully depolarized and no further depolarization by positive electric field could occur. At 0.02 mW/mm^2^ irradiance and 10 ms pulse width, the maximum voltage drop across the degenerate photoreceptor cell body was -1.6 mV (Figure 7D). These findings indicate that retinal activation in the dark at threshold (0.014 mW/mm^2^ at 10ms), which is lower than that required for direct stimulation of bipolar cells (0.067 mW/mm^2^) occurs due to presence of residual photoreceptors near the negative potential from the peripheral return electrode of monopolar implants (red curve in Figure 6A). In light-adapted conditions, however, bipolar cells are depolarized due to the influence of residual photoreceptors outside the implant, and their stimulation threshold decreases to 0.024 mW/mm^2^ at 10 ms (black curve in Figure 6A). Chronaxie of the S-D curve in light-adapted conditions (3.2 ms) indicates stimulation of bipolar cells rather than direct activation of PRs.

With bipolar photovoltaic arrays, much tighter confinement of electric field caused by return electrodes around each pixel (Figure 1B) resulted in much smaller hyperpolarization of healthy photoreceptors outside the device at stimulation threshold (Figure 7A, blue dashed) - a maximum of -0.5 mV at 0.20 mW/mm^2^. This is approximately a third of the value observed with monopolar implants at their threshold (0.020 mW/mm^2^) (Figure 7A, blue asterisks). Degenerate photoreceptors polarize in the electric field from bipolar arrays even less (Figure 7A, red dashed). Distributed return electrodes in bipolar arrays provide a tight field confinement which allows avoiding the activation of photoreceptors adjacent to the implant. However, with a very non-uniform illumination, a beam shifted to the edge of such implant may still induce direct stimulation of photoreceptors in the dark, as shown in Figure 5 (purple, dashed).

## 4. Discussion

### 4.1. Surrounding photoreceptors affect the electrical stimulation threshold

One of the challenges with bipolar photovoltaic arrays (PRIMA) is that electric field is so constrained by the local return electrodes that with pixels much smaller than 100 μm, it cannot reach the bipolar cells in the INL [3]. This issue could be mitigated using an array of monopolar pixels. Such devices provide deeper penetrating electric field, albeit with reduced contrast [10]. However, this study identified a potential problem: in the dark, photoreceptors can be stimulated directly with 5-10 ms pulses by negative potential generated by a common return electrode near the edge of the implant. This suggests that if the device is placed at the border of a scotoma [2], healthy photoreceptors nearby may skew the percepts near the implant’s edge.

Photovoltaic prosthesis PRIMA, currently used in clinical trials, has bipolar pixels and therefore much smaller negative potential near its edges than that with monopolar arrays (Figure 6C, D). Therefore, it is not expected to stimulate the adjacent photoreceptors under uniform illumination, and PRIMA patients indeed did not report perceptions of a bright line along the edge of the implant. Future implant designs should therefore be bipolar, with better penetration of electric field into the retina achieved with 3-D electrodes [26] rather than with the field summation from monopolar pixels [10].

Another effect of the remnant photoreceptors surrounding the implant may influence the threshold of bipolar cells near the edge of the device even with bipolar arrays – depolarization of ON BCs under ambient lighting. To avoid the potential influence of remaining photoreceptors on prosthetic vision, implant should be placed at least 150 µm from the healthy retina.

The stimulation threshold of photoreceptors drops with increasing pulse width, with no observable saturation in the range of our studies – up to 10 ms. Our previous studies demonstrated a similar trend continuing at least to 100 ms [15]. Other studies have also reported that subretinal stimulation of degenerate photoreceptors can elicit retinal responses [27]. Incredibly low stimulation thresholds of photoreceptors at long exposures may explain the low irradiance thresholds observed with other subretinal optoelectronic implants [28].

### 4.2. Potential mechanisms of signal propagation and associated thresholds

Under photopic conditions, the recorded threshold for monopolar implants in LE rats was about 3-4 times lower at all pulse widths compared to that with L-AP4 injection (Figure 6A, black vs. green). Surprisingly, under dark-adapted conditions, stimulation threshold increased for pulse durations <8ms (Figure 6A, red) but decreased for longer pulses. It indicates that stimulation in scotopic conditions may be driven by a combination of photoreceptors and bipolar cells. Photoreceptors are the most sensitive in the dark, when they maintain a membrane potential of approximately -40 mV [29], resulting in ON bipolar cells maintaining a membrane potential of - 41 mV [30], about 2.5 mV below the synaptic threshold of ON bipolar cells to release glutamate onto ON RGCs, when calcium channels open at -38.5 mV [13]. Similar state occurs with L-AP4 (mGluR6 agonist) injection. Under photopic conditions, the ON bipolar cells are depolarized up to 10 mV by photoreceptors outside the implant and hence exhibit lower electric threshold (black S-D curve in Figure 6A). Dark-adapted thresholds reflect a transition from bipolar cells-mediated stimulation with shorter pulses (<1 ms) to photoreceptors-mediated response with longer pulses (Figure 6A, red). L-AP4 injection (green line) reveals the threshold for stimulation of bipolar cells alone, without any contribution from photoreceptors.

Bipolar cells’ threshold could be influenced by the remaining photoreceptors surrounding the implant though either the AII amacrine network [31] or through direct input from degenerated PRs (PRs lacking outer segments) above the edge of the device. As rod bipolar cells directly communicate to AII cells, they could laterally transmit signals from outside the implant to bipolar cells above the implant, thereby affecting their threshold (Figure 8A) within the reach of AII cells (100μm) [32]. Degenerated PRs can also receive signals from adjacent healthy PRs through gap junctions located in the cone pedicles and rod spherules [33, 34], which are still preserved in local degeneration. This way, degenerate PRs can be hyperpolarized by illuminated healthy photoreceptors and affect the membrane potential of ON BCs near the edge (Figure 8B) resulting in reduced thresholds. Direct input from healthy PRs to BCs above the implant is unlikely as rod bipolar cells have an average receptive field of 67 μm, and cone bipolars have an average receptive field of 43 μm [35], which is less than the distance from the device edge to its photovoltaic pixels separated by the 250 μm wide common return electrode in monopolar arrays.

**Figure 8.**
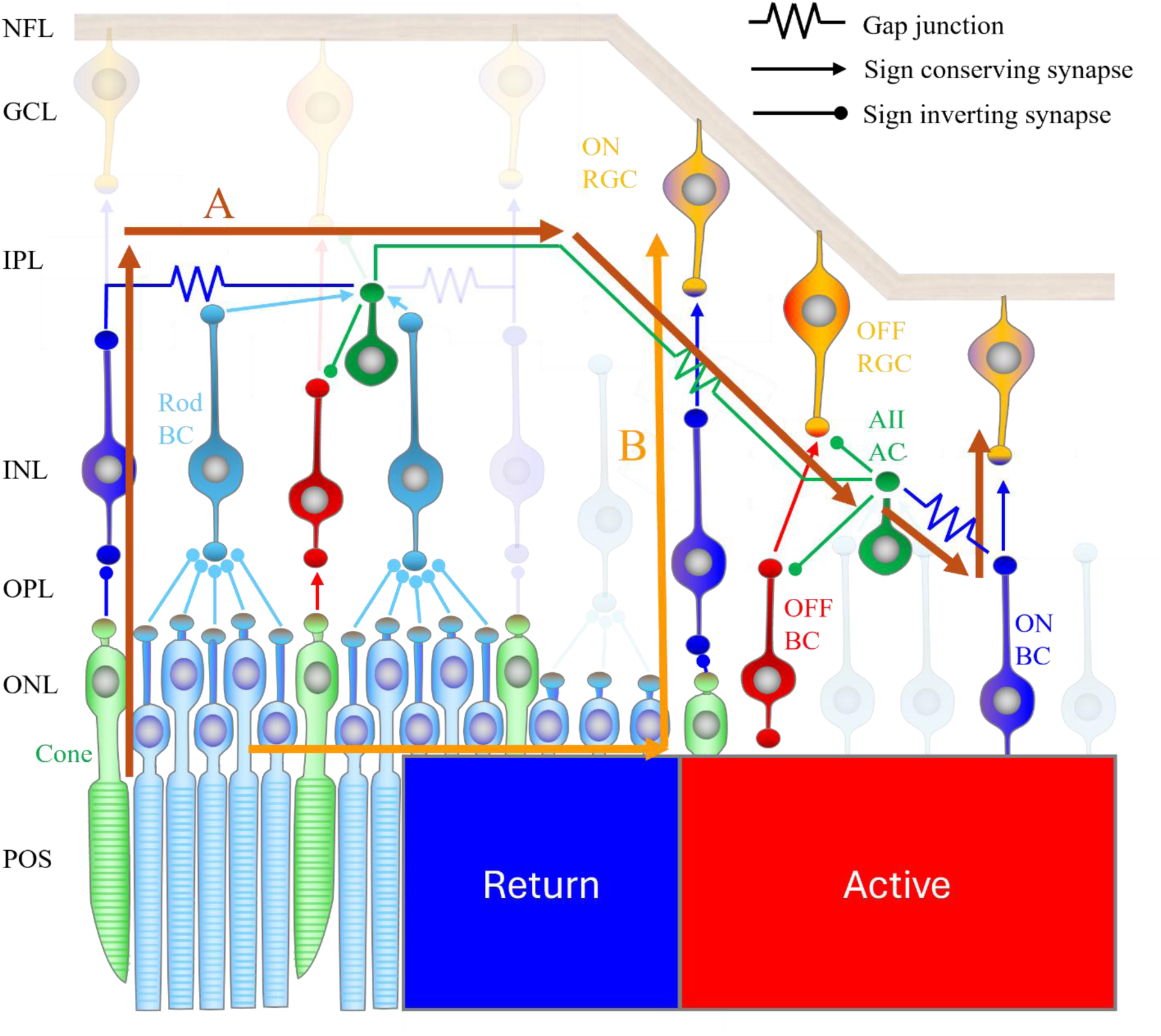
Possible mechanisms to how dark adaptation of photoreceptors around the implant can influence the threshold of the inner retina at the center of the implant. **A.** Rod photoreceptors (light blue) send signals to rod BCs which then depolarize AII ACs (dark green). ACs form a network of coupled cells in the inner plexiform layer which allow signals to spatially disperse laterally, resulting in a change in membrane potential of BCs and RGCs further than the PRs original location. Additionally, cones (light green) are connected to ON BCs (dark blue) which are electrically coupled to AII ACs. **B.** Rod and cone PRs form a network of electrically coupled neurons which can transmit signals laterally resulting in degenerate PRs reducing the threshold of ON BCs. These mechanisms of lateral communication allows remaining PRs to influence the response of BCs directly under active electrodes.

Additionally, we found that strong bleaching (over-hyperpolarizing) of PRs decreases the VEP amplitude in response to electrical stimulation (Figure 4B) which is consistent with previous findings [21, 22]. PRs strongly hyperpolarized by bleaching cannot be hyperpolarized much further by negative electric field near the common return electrode. Similarly, ON BCs strongly depolarized under bleaching cannot be stimulated much further by positive electric field, thus reducing the VEP response at the same irradiance. This suggests that strong surrounding natural illumination can attenuate prosthetic vision by decreasing the retinal response, whilst a weak illumination may reduce the stimulation threshold.

### 4.3. Different thresholds with monopolar and bipolar arrays

Eliminating the effect of photoreceptors by L-AP4 injection in LE rats or using RCS rats, demonstrates a major difference between the stimulation thresholds of bipolar cells with monopolar and bipolar arrays (0.06 mW/mm^2^ vs. 0.20 mW/mm^2^ at 10 ms, Figure 6A, B). Voltage drop across bipolar cells at their respective thresholds is shown in Figures 6C-F. As VEP is a population response and RGC response is the weighted sum of their input from BCs above threshold (in this case a weighted sum of the positive step of potential across BCs), we also calculated the average voltage across BCs. For bipolar array, where there are negative areas in each pixel, only positive voltages (which depolarize BCs) were used to calculate the average. At the threshold irradiances, the average potential drop across BCs over the illuminated monopolar array was 4.6 mV (Figure 6E, green), over bipolar array it was 4.7 mV (Figure 6F, red). The peak voltage with bipolar arrays at 10 ms pulse width was 11.8 mV (Figure 6F), consistent with our previous findings of approximately 11 mV [3].

A much lower stimulation threshold at photopic conditions than with L-AP4 for monopolar arrays (0.024 mW/mm^2^ vs. 0.067 mW/mm^2^ at 10ms, Figure 6A) can be explained by increased resting potential of ON BCs at the edge of the implant due to photoreceptors input [25]. As discussed above, the maximum lateral range of photoreceptors’ influence on BCs in the rat retina is 90 µm [36], which is equivalent to 4.5 monopolar pixels of 20 μm, or less than one 100 μm bipolar pixel. At this irradiance, BCs directly above the active monopolar pixels on the edge experience an average 1.6 mV drop (Figure 6E, black), which is sufficient to elicit a response, whilst a single 100 µm pixel on the edge of the bipolar array at its stimulation threshold (0.20 mW/mm^2^) has an average potential drop of 1.4 mV across BCs (Figure 6F). Although these values are relatively similar, at this irradiance, the voltage drop across bipolar cells in the center of the bipolar array experience a much larger drop in electric field (4.7 mV in bipolar array vs 1.8 mV in monopolar array).

This effect was not observed with bipolar arrays (Figure 6B) because of non-uniformity of illumination in our setup. Electric field above bipolar arrays replicate the illumination pattern much better than monopolar arrays, which smear it due to strong cross-talk between electrodes (Figure 6 C, E vs. D, F). However, when the beam center was shifted to the edge of the bipolar implant, stimulation threshold decreased similarly to that with monopolar arrays (Figure 5, blue).

In human retina, rod bipolar cells and AII amacrine cells have a dendritic field diameter of 109.2 ± 68.4 µm and 47.3 ± 25.4 µm, respectively [37], suggesting that a single rod can affect bipolar cells (rod → rod bipolar cell → AII amacrine cell → ON bipolar cell) at distances of up to 78.3 ± 46.9 µm. Therefore, placing the implant at least 150 µm from the edge of the scotoma should avoid PRs influence on stimulation threshold of the implant.

Modelling also demonstrates that photoreceptors can be stimulated by the large return electrodes of monopolar arrays, but not with bipolar implants (Figure 7A). Although the peak negative potential above monopolar and bipolar arrays is similar (-1.3 mV vs. -1.5 mV, Figure 6D), higher negative potential is concentrated near the edge of the monopolar implant (Figure 6C), while in bipolar arrays it is distributed over narrow local returns surrounding each pixel (Figure 6D). Assuming 70 μm long healthy photoreceptors at the edge, the average voltage drop across PR over a 100 μm wide area is -1.2 mV for monopolar and -0.5 mV for bipolar arrays. As a result, with monopolar arrays, the PR threshold is 3 times below that of BCs (0.02 vs. 0.06 mW/mm^2^ at 10ms), whilst with bipolar arrays, the BCs threshold (0.2 mW/mm^2^) is reached before the PR threshold, as summarized in Table 1.

**Table 1.** Average voltage (averaged spatially across 100 µm) across PRs at the edge of the return (from 0 µm and 70 µm), BCs on the edge of the active (from 0 µm and 50 µm), and BCs on the center of the active electrodes. Monopolar implants first reach PR threshold of -1.2 mV across the cell at 0.015 mW/mm^2^, before reaching PR influenced BC threshold of 1.6 mV across the cell at 0.024 mW/mm^2^. Uninfluenced BCs at the center of the implant have a threshold almost 3 times, with 4.6 mV drop across the cell at 0.062 mW/mm^2^. PRIMA reaches uninfluenced BC threshold first at 0.2 mW/mm^2^. Thresholds for each cell are shown at the bottom of the table. Voltages at or above thresholds are shown in bold.

|  | Irradiance<br>(mW/mm<sup>2</sup>) | PRs<br>(mV) | BCs (Edge)<br>(mV) | BCs (Center)<br>(mV) |
| --- | --- | --- | --- | --- |
| MP | 0.015 | **-1.2** | 0.7 | 1.13 |
|  | 0.024 | **-1.6** | **1.6** | 1.8 |
|  | 0.062 | **-4.2** | **2.8** | **4.6** |
| PRIMA | 0.200 | -0.5 | 1.4 | **4.7** |
| Thresholds | - | -1.2 | 1.6 | 4.6 |

### 4.4. Dominance of the ON over OFF pathway

Signaling mechanisms, including the effect of dark adaptation and of L-AP4 blocker discussed above, assume that the ON pathway dominates the retinal response to subretinal photovoltaic stimulation, which may explain why patients report only bright phosphenes [38], as opposed to both bright and dark percepts reported with cortical [39, 40] and epiretinal stimulation. For the OFF pathway, shifts of thresholds upon dark adaptation and L-AP4 injection are expected to be the opposite of that observed in this study. ON bipolar cells are hyperpolarized under L-AP4 due to maximum activation of metabotropic glutamate receptors, resulting in an increased threshold, whilst OFF bipolar cells are not directly influenced by L-AP4. Additionally, the hyperpolarization of ON and rod bipolar cells results in the suppression of the amacrine cell network, thereby reducing the glycinergic rectification of the OFF pathway [41]. Therefore, dark adaptation or L-AP4 application is expected to reduce the threshold for OFF bipolar cells – opposite of the effect observed in our study.

Dominance of the ON pathway may be explained by several factors. Since rod BCs have only the ON type, rod-dominated rat retina (99%) is naturally dominated by the ON pathway [42]. An additional factor could be that much lower spontaneous firing rate in ON than in OFF RGCs in the degenerate retina [43] results in a higher signal-to-noise ratio in ON vs. OFF pathway, making it perceptually dominant. Yet another reason could be the rectification of the OFF pathway by AII amacrine cells [41], resulting in the ON bipolar responses dominating the retinal output [41]. Additionally, stimulation threshold of ON BCs may be lower than OFF BCs because ON BCs are depolarized whilst OFF BCs are hyperpolarized in the degenerate retina due to the absence of glutamate input from PRs. ON BCs are also longer than OFF BCs, resulting in a higher voltage drop across the cell causing more depolarization in ON BCs than OFF BCs [44].

The ON pathway dominance has been previously reported in both subretinal electrical stimulation [45] as well as in BC targeted optogenetics [46]. It has been described that blocking glycinergic inhibitory inputs to OFF BCs and OFF RGCs from AII amacrine cells results in 50% reduced rectification [45], suggesting that coupling of the rod BC to AII AC network plays a significant role in causing ON pathway dominance. Additionally, in BC-targeted optogenetic therapies [46], ON responses dominate (68.7% ON vs. 25.7% OFF) in the treated *rd1* mouse retina, which also indicates the role of AII amacrine cells in shaping the ON pathway dominance.

### 4.5. Effect of synaptic plasticity on stimulation thresholds

Interestingly, thresholds in LE rats (local degeneration) with L-AP4 injection and RCS rats (full degeneration) [4] are the same, even though the baseline membrane potentials are different [25]. L-AP4, as a glutamate agonist (equivalent to maximum glutamate concentration or dark conditions), causes hyperpolarization of ON bipolar cells, whilst the loss of the glutamate input due to missing photoreceptors in the degenerate RCS retina results in depolarization of ON bipolar cells (equivalent to bleached retina under bright light conditions) [25]. It was shown previously that despite the loss of glutamate input to ON BCs, the ON RGCs maintain the same spontaneous firing rate throughout the course of retinal degeneration [43], indicating adaptation of the retina by reducing the ON BC synaptic output (synaptic plasticity) [47].

This reduction in synaptic output to RGCs has been previously reported in *rd1* and *rd10* mouse retina [48]. It was shown that although the number of ribbon synapses did not change in retinal degeneration, excitatory post-synaptic potentials (EPSCs) in ON RGCs were reduced in the degenerate retina because of the loss of glutamate input in ON BCs from photoreceptors (Figure 9A). As a result, the lack of glutamate input to BCs in RCS rats, and full activation of the glutamate receptor in BCs with L-AP4 in LE rats yield the same threshold for bipolar cell activation (Figure 9B). Such “self-calibration” of the inner retina by synaptic plasticity preserves the same perceptual thresholds between healthy and degenerate retina and plays a crucial role in maintaining the same stimulation thresholds.

**Figure 9.**
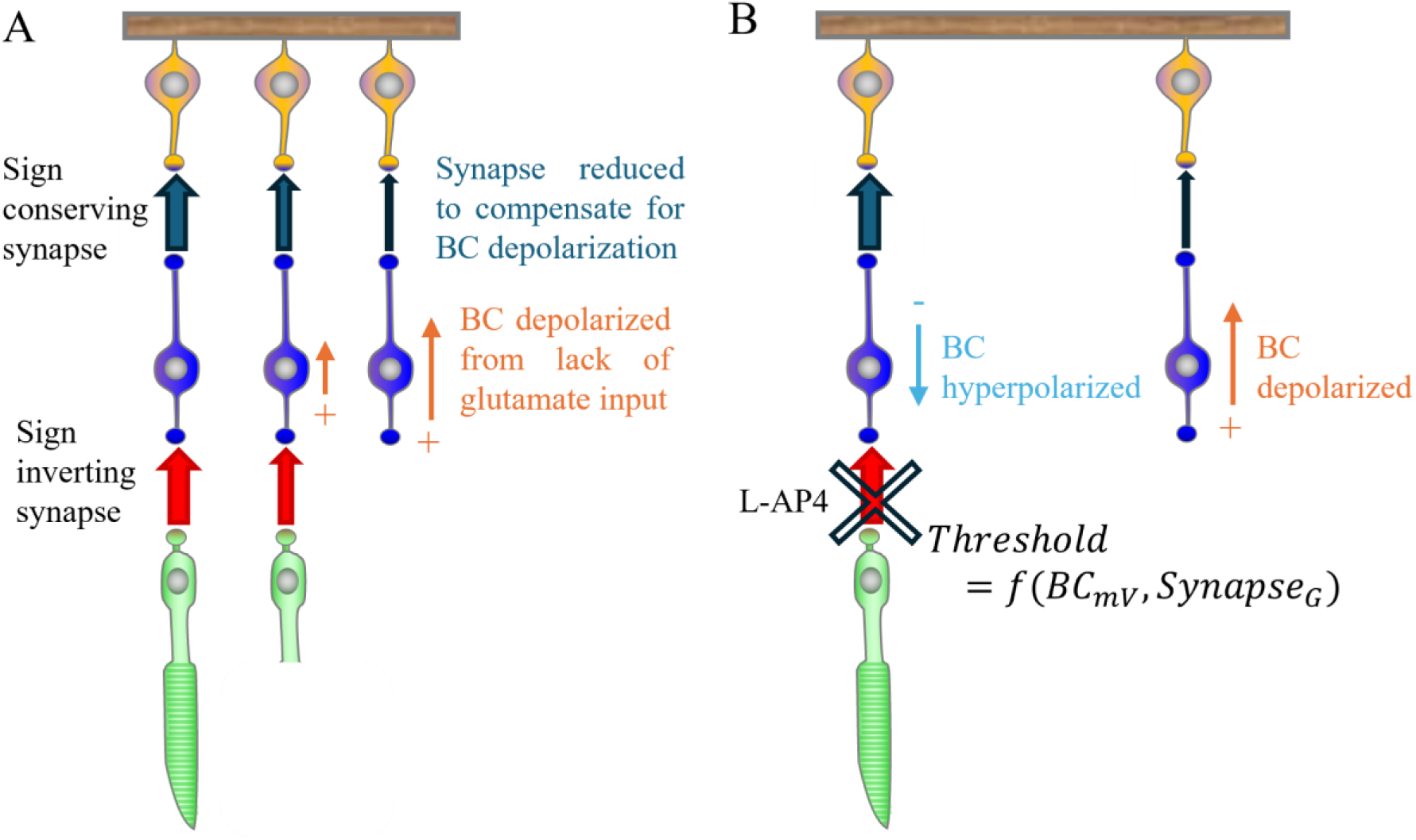
Synaptic plasticity in the degenerate retina preserves stimulation thresholds. **A.** Reduction of synaptic conductance between ON BCs and ON RGCs compensates for the depolarization of BCs due to lack of glutamate input from PR degeneration. **B.** Injection of mGluR6 agonist L-AP4 causes ON BCs to be hyperpolarized, increasing the threshold to 0.06 mW/mm^2^. In the degenerate retina, ON BCs are depolarized which suggests a reduced threshold but as the synapse between ON BC and ON RGC is reduced to compensate for this, the threshold results in same value of 0.06 mW/mm^2^.

## 5. Conclusions

Some photoreceptors are often retained during retinal degeneration, especially in maculopathies. Our study indicates two different effects by which photoreceptors influence the subretinal electrical stimulation: 1) if exposed to negative electric potential, photoreceptors may respond at significantly lower threshold with longer pulses (∼10ms) compared to direct stimulation of bipolar cells above the implant, and 2) bipolar cells near the edge of the implant are more depolarized due to photoreceptor input in photopic conditions, resulting in a reduced threshold. To avoid a bright percept near the edge of the implant, devices should be designed such that they do not generate a negative potential spilling beyond their edges. As such, bipolar arrays, having local return electrodes in each pixel provide a more promising design for subretinal prosthesis. For small bipolar pixels, electric field penetration into the retina with flat arrays is insufficient [49], and 3-D electrodes are required to provide better penetration of electric field into the INL, while retaining good lateral confinement [50]. But even with bipolar arrays, to avoid the depolarizing effect of residual photoreceptors on bipolar cells, implants should be placed at least 150 µm from the edge of the scotoma.

## Acknowledgements

Studies were supported by the National Institutes of Health (Grants R01-EY- 035227, P30-EY-026877), the Department of Defense (Grant W81XWH-2210933), and AFOSR (Grant FA9550-19-1-0402). We would like to thank Stephen A. Baccus (Stanford University, CA) for insightful discussions on neural encoding in the healthy and degenerate retina.

